# Analysis of the surface topology of respiratory syncytial virus particles that form on the surface of virus-infected cells

**DOI:** 10.1101/2025.07.26.666719

**Authors:** Chris E. Jeffree, Thong Beng Lu, Boon-Huan Tan, Richard J. Sugrue

## Abstract

The surface topology of virus filaments on respiratory syncytial virus (RSV)-infected cells was examined using field emission gun-scanning electron microscopy (FEG-SEM) and atomic force microscopy (AFM). FEG-SEM analysis of the surface of RSV-infected cells labelled with an anti-G protein antibody revealed the presence of virus filaments and clusters of the G protein distributed intermittently along their surface. RSV-infected cells thinly coated with chromium were imaged using FEG-SEM and revealed a distinct structured surface topology consisting of closely packed surface domains. The G protein clusters were only associated within a subset of these domains which suggested that this structured topology was mainly derived from the host cell, and the presence of the cell glycocalyx that coats the virus filaments was further suggested. Imaging of RSV-infected cells using AFM was undertaken as a different but complementary approach to the FEG-SEM analysis. Imaging using AFM revealed a similar structured surface topology on the virus filaments to that observed in the FEG-SEM analysis, indicating the consistency in the appearance of the virus surface topology using these different methods. Collectively, this study provides the first detailed imaging of the surface topology of the virus filaments as they form on RSV-infected cells. The imaging data is consistent with the envelopment of the virus filaments by the glycocalyx and highlights the complexity of the spatial organisation within the viral envelope.

## Introduction

Respiratory syncytial virus (RSV) is the major cause of severe lower respiratory tract disease in several high-risk groups which include young children, the elderly and immunocompromised adults. On the surface of RSV-infected cells the mature infectious RSV virus particles assemble as cell-associated membrane-bound filamentous structures [1, 2]. We refer to these as virus filaments, and on permissive cell models these virus filaments have been demonstrated to play a role in facilitating virus transmission, and treatments that prevent virus filament formation led to reduced levels of virus infectivity[3, 4]. Virus filaments are also detected on organoid cell systems infected with RSV [5] providing evidence that these structures have a physiological and clinical relevance. Although the structural organisation of the virus proteins within mature RSV particle has been largely characterised [6], the virus assembly mechanism has not yet been completely defined. However, the involvement of several host-cell factors has been implicated in the RSV particle assembly process, including the cortical F-actin network and the associated cell signalling pathways [4, 7–10].

RSV assembly occurs within specialized lipid raft microdomains on the cell surface [11–13], and a role for specialized lipid-raft domains called caveolae is suggested[14, 15]. The virus filaments are therefore bounded by a lipid envelope that is derived from the host cell at the site of virus assembly. Within the virus envelope three virus-encoded integral membrane glycoproteins are inserted, the fusion (F) protein, the attachment (G) protein and the Small Hydrophobic (SH) protein. The G protein is established as the RSV cell-attachment protein [16, 17], and the F protein initiates fusion of the virus envelope and cell membrane during virus entry into the host cell[18], and the interaction between the F and G proteins on the surface of the virus filaments mediates this localised virus transmission [19–21]. The function of the SH protein in the virus particle is still currently uncertain but it may play a role in mitigating the host response to infection[22]. The virus-expressed Matrix (M) protein is a peripheral membrane protein that interacts with the virus envelope and forms a protein layer beneath the virus envelope. The nucleocapsid (NC) core is formed by the interaction of the viral genomic RNA (vRNA) with the nucleocapsid (N) protein, the phosphoprotein (P), the M2-1 protein and the large (L) protein [23–27]. In RSV-infected cells the NC accumulate in the cytoplasm within inclusion bodies and the NCs are trafficked from the inclusion bodies to the site of RSV assembly where they are packaged into the interior of the virus filaments [28, 29].

Since virus filaments play a direct role in facilitating RSV transmission, a better understanding of how these structures form should enable the development of novel antivirus strategies. This includes an improved understanding of the organization of the components that are displayed on the surface of the virus filaments i.e. the surface topology of the virus filaments. The surfaces of the virus-infected cells are surrounded by a thin polysaccharide matrix coating structure called the glycocalyx. This consists of specific glycolipids and membrane-bound glycoproteins, and the glycocalyx is associated with several important cellular functions e.g. defining plasma membrane shape [30]. During RSV particle assembly the virus filaments become enveloped by the glycocalyx [31], and the presence of the glycocalyx on the surface of the virus filaments suggests an additional level of complexity on the surface topology of the virus envelope. In this context, previous detailed ultrastructural analyses of RSV-infected cells using transmission electron microscopy have reported that the surface of the virus filaments were coated by a relatively thin amorphous layer into which the virus glycoproteins were embedded [32–34]. These earlier studies did not address the origin of this amorphous layer on the surface of the virus filaments, but in light of recent evidence that cell glycocalyx coats the virus filaments during virus particle assembly, we presume that the thin amorphous layer that was observed in these earlier studies indicated the presence of the glycocalyx. To examine this further we have employed high-resolution imaging of the surface of virus infected cells. We have previously used scanning electron microscopy (SEM) to examine the virus filaments that form on the surface of RSV-infected [2]. In this current study has extended our earlier work to examine the complexity of the surface topology of the virus envelope in greater detail using both SEM and in addition atomic force microscopy (AFM). This current experimental approach is the first study to highlight the complexity of the surface topology of the virus filaments as they form during virus particle assembly on the surface of virus-infected cells.

## Results and discussion

### 1. Detection of anti-G-stained virus filaments on the surface of virus infected cells using field emission gun scanning electron microscopy (FEG-SEM)

We used immuno-scanning electron microscopy (ISEM) to examine the virus filaments that form on the surface of A549 cells infected with RSV. The G protein is a major virus component that is abundantly displayed on the surface envelope along the entire length of the virus filaments and therefore demarcates these structures [19]. In addition, the G protein is embedded within the glycocalyx that coats the virus filaments [31] and can therefore be used as a marker to locate the virus-associated cell glycocalyx. The antibody anti-G (that detects the G protein) is also compatible with the procedures used to prepare virus-infected cells for use in scanning electron microscopy analysis [2], and when appropriate anti-G staining was also used to confirm infection and the presence of virus filaments. We initially applied a 20-nm thick conductive coating of carbon by vacuum evaporation to prevent charging of the biological specimens in the electron beam in the scanning electron microscope. In the ISEM analysis the cells were labelled with anti-G, and the presence of anti-G labelling was detected using the secondary antibody anti-mouse IgG conjugated to colloidal gold particles of uniform size (either 5 or 10nm diameter), and the antibody-labelled cells examined using a Field Emission Gun– Scanning Electron Microscope (FEG-SEM). In the FEG-SEM analysis the cells in the same field of view were imaged simultaneously using two different detectors. A secondary electron (SE) detector that allows visualisation of the surface topology of the cells and imaging of the virus filaments. An Yttrium Aluminium Garnet (YAG) backscattered electron (BSE) detector was configured to detect the strong backscattered electron signal produced by colloidal gold and this enabled us to detect the presence of the bound secondary antibody that is conjugated to colloidal gold. In the images obtained using the YAG BSE detector the colloidal gold appears as white spots indicating the presence and location of the bound anti-G antibody and hence the distribution of the G protein. Osmium tetroxide (OsO_4_) was used in the specimen preparation to preserve cellular ultrastructure, and the spectral characteristics of elemental osmium (Os) is close to that of elemental gold. Therefore, in the same images obtained using the YAG BSE detector, the strong bright spots from the gold particles are detected, together with a low-level of general background signal due to the presence of the OsO_4_. This faint background signal delineates the cell topology and the virus filaments and facilitates the alignment of the antibody signal on the surface of the virus filaments by cross-referencing the images taken with the YAG backscatter and SE detectors of the same field of view. By comparing the images obtained in the same field of view using both detectors the distribution of the G protein and the virus filaments on the RSV-infected cells can be examined.

Cells were mock-infected and RSV-infected and at 20 hrs post-infection (hpi) the cells were stained with anti-G and examined by immunofluorescence (IF) microscopy. The presence of anti-G labelling was detected using anti-mouse IgG conjugated to the fluorescence Alexa probe AL488. Fluorescence staining of cells was only detected in cells infected with RSV and was absent in mock-infected cells (Fig. 1A), confirming both virus infection and the specificity of the anti-G antibody.

**Figure 1.**
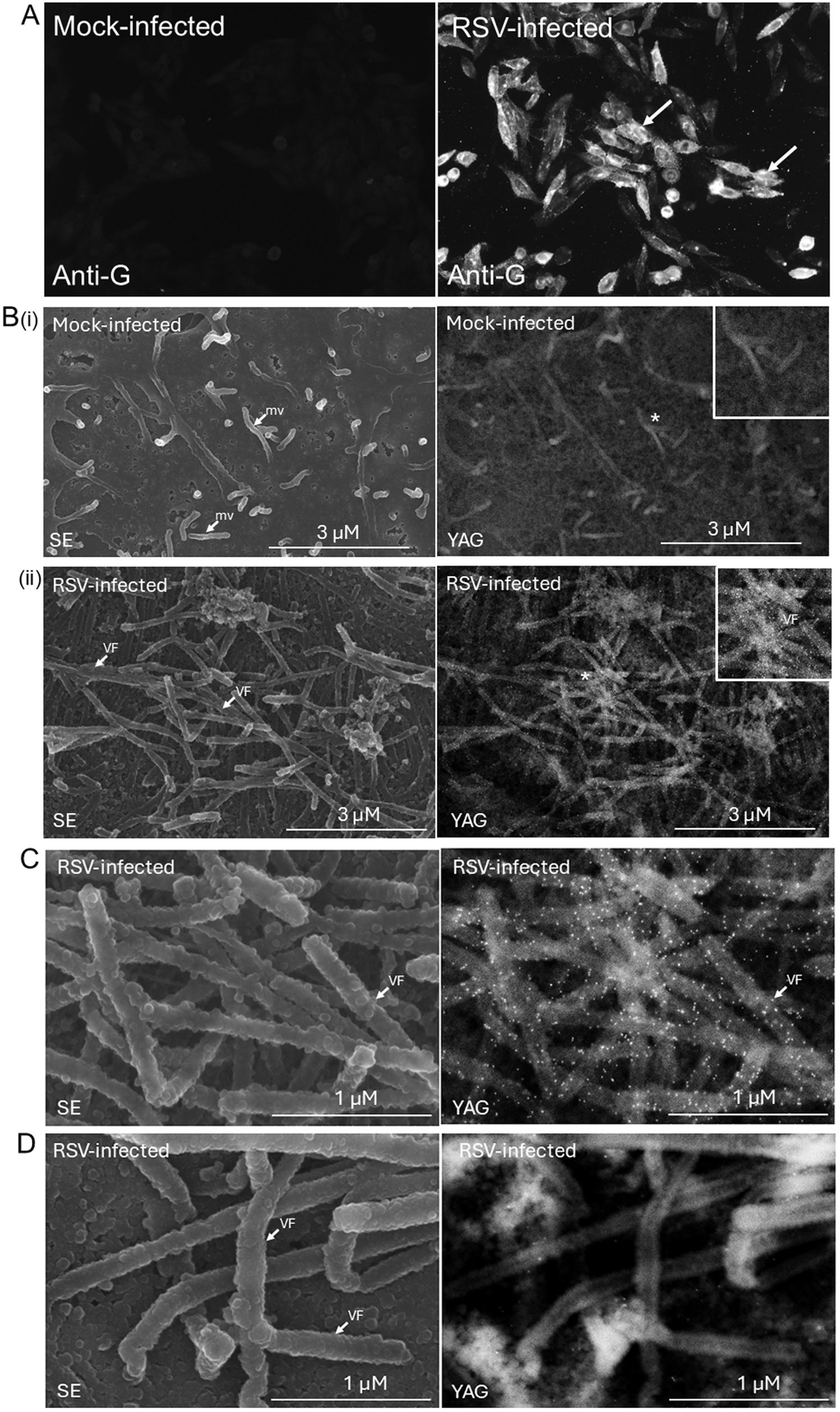
Localization of the G protein on RSV filaments on the surface of virus-infected cells using immune-scanning electron microscopy. (A) At 18 hrs-post infection (hpi) mock-infected and RSV-infected A549 cells were incubated with anti-G and anti-mouse AL488 and the bound antibody was detected using immunofluorescence microscopy (objective x40 magnification). The virus-infected cells are highlighted (white arrows). (B) At 18hpi (i) Mock-infected and (ii) RSV-infected A549 cells were incubated with anti-G and anti-mouse IgG conjugated to 10-nm colloidal gold. The surface of the monolayers was visualized in the FEG-SEM using the secondary electron (SE) and YAG BSE detector (YAG) as indicated. The presence of gold particles in the YAG BSE image are presented as small bright points. The presence of the microvilli (mv) and virus filaments (VF) are highlighted in the SE image, and the insets in the YAG BSE image are enlarged regions of the area in the main indicated by *. (magnification x15,000). The RSV-infected cells were labelled with (C) anti-G and anti-mouse IgG conjugated to 10-nm colloidal gold and (D) anti-SH and anti-mouse IgG conjugated to 10-nm colloidal gold imaged using the SE and YAG BSE (YAG) detectors as indicated (magnification x50,000). The virus filaments (VF) are highlighted.

Cells were mock-infected and RSV-infected and at 20hpi the cells were labelled with anti-G and anti-mouse IgG conjugated to 10nm colloidal gold particles (anti-mouse IgG-10nmAu). Analysis of the antibody labelled mock-infected and RSV-infected cells using the SE detector confirmed the integrity of the cell surface (Fig. 1B(i) and (ii)). Analysis of the surface of mock-infected cells using the SE detector (Fig. 1B(i)) showed the presence of numerous relatively small cell surface projections prior to virus infection, such as microvilli that were approximately 300-500nm in length. The YAG BSE detector indicated the absence of colloidal gold on surface of these cells, indicating with the absence of anti-G staining of the mock-infected cells and consistent with the IF microscopy imaging. In contrast, on the surface of the virus-infected cells the presence of long virus filaments (>1μm in length) was detected using the SE detector (Fig. 1B(ii)), being significantly longer than the microvilli and consistent with previous observations [1, 2]. The presence of anti-G staining on these cells was detected by the YAG BSE detector and was localised on the virus filaments. This was consistent with the identification of these anti-G-stained filamentous projections as the virus filaments. Analysis of the RSV-infected cells using the FEG-SEM at higher magnification using the SE detector more clearly defined the virus filaments and the images suggested a structured surface topology on the virus filaments (Fig. 1C). In the YAG BSE detector the anti-G labelling along the entire length of the virus filaments on these cells was observed (up to 3 gold particles per cluster). Although a localised clustering of the anti-G labelling at locations along the surface of the virus filaments was noted, the clustered anti-G labelling was intermittently distributed along the surface of the virus filaments (Fig. 1C). We also failed to detect the presence of the abundant and shorter microvilli on the surface of the virus-infected cells, suggesting a reduction in the numbers of microvilli following virus infection. Since microvilli are stabilised by the cortical F-actin network, this observation was consistent with the F-actin remodelling that has been reported in RSV-infected cells[4, 7, 8, 10] and which would be expected to lead to changes in the structure of the microvilli. In this context, in cells expressing the recombinant G protein the recombinant G protein is transported into filamentous projections that resemble the microvilli on the cell surface [8, 31].

To confirm the specificity of the anti-G binding on the virus filaments, as a negative control the infected cells were also labelled with anti-SH and anti-mouse IgG-10nmAu (Fig. 1D). The anti-SH antibody detects an antibody binding epitope on the cytoplasmic tail of the SH protein [35] and this antibody binding epitope is not expected to be displayed either on the intact surface of virus-infected cells nor on the surface envelope of the structurally intact virus filaments. While virus filament formation was detected using the SE detector, antibody labelling on the surface of the virus filaments was not observed by the YAG BSE detector (Fig. 1D). This confirmed the binding specificity of the anti-G antibody labelling on the virus filaments and that any non-specific binding of the anti-mouse IgG-10nmAu to the cell surface was below the level of detection.

Although virus filaments are the predominant form of the virus that is detected upon RSV-infected cells, at the later stages of infection a proportion of these filamentous virus particles become detached from the infected cell and these cell-free virus particles adopt a pleomorphic morphology. It currently unclear if this transition to a cell-free virus pleomorphic morphology is a non-specific process that arises following mechanical detachment of the virus filaments from the cell, or if it is mediated by a specific molecular mechanism. Although the precise mechanism that underpins this transition is not completely understood, these cell-free virus particles are expected to play a role in facilitating horizontal virus transmission. However, conditions that inhibit virus filament formation also reduce the levels of cell-free virus infectivity, suggesting that virus filament formation is a prerequisite for the production of the cell-free virus infectivity [3]. The cell-free virus particles present in the virus inoculum also adopt the pleiomorphic morphology, and to more clearly distinguish the virus particles in the virus inoculum from the virus particles that form during virus assembly we also used ISEM to examine the surface of virus-infected cells at the early and lates stages of infection. During the early stages of virus infection (prior to virus particle assembly) this pleiomorphic virus particle morphology would be expected to be present on the surface of the cells. This virus particle morphology would be distinct from the virus filaments that form on the surface of infected cells at the later stages of infection following virus particle assembly.

The ISEM analysis was therefore used to enable a clear distinction to be made of the two different types of virus particle morphologies that are present on the cell surface during the different stages of RSV infection. In this analysis 2hpi and 30 hpi were used to represent early and late time points of infection respectively. Cells were infected with RSV and at 2 and 30 hpi the cells were stained using anti-G and anti-mouse IgG conjugated to AL488 and the stained cells examined using a confocal microscopy at a focal plan that allowed imaging of the virus particles (Fig. 2A). At 2 hpi a prominent punctate anti-G staining pattern was apparent which was consistent with the cell-free virus in the inoculum attaching to the cell surface (Fig. 2A(i)), while at 30 hpi the prominent filamentous anti-G staining pattern was observed and was consistent with the formation of virus filaments (Fig. 2A(ii)). The cells were infected with RSV and at 2 hpi (Fig. 2B) and 30hpi (Fig. 2C) and labelled with anti-G and anti-mouse IgG-10nmAu and imaged by FEG-SEM using the SE and YAG backscatter detectors. At 2 hpi only pleiomorphic-shaped anti-G-stained particles on the surface of these cells were present and was consistent with the attachment of pleomorphic shaped virus particles on the surface of the cells at the initial stages of infection (Fig. 2B(i)). These ranged from virus particles that exhibited a spherical morphology (approximately 300 nm in diameter) to virus particles that displayed both partial spherical and filamentous morphologies (Fig. 2B(ii)). At 30 hpi only abundant quantities of the anti-G-stained virus filaments could be detected on the surface of the infected cells using the SE and YAG backscatter detectors (Fig. 2C(i)). The anti-G staining on the virus filaments was apparent along their length, and the presence of clusters of colloidal gold particles suggested a localised concentration of the G protein at these locations.

**Figure 2.**
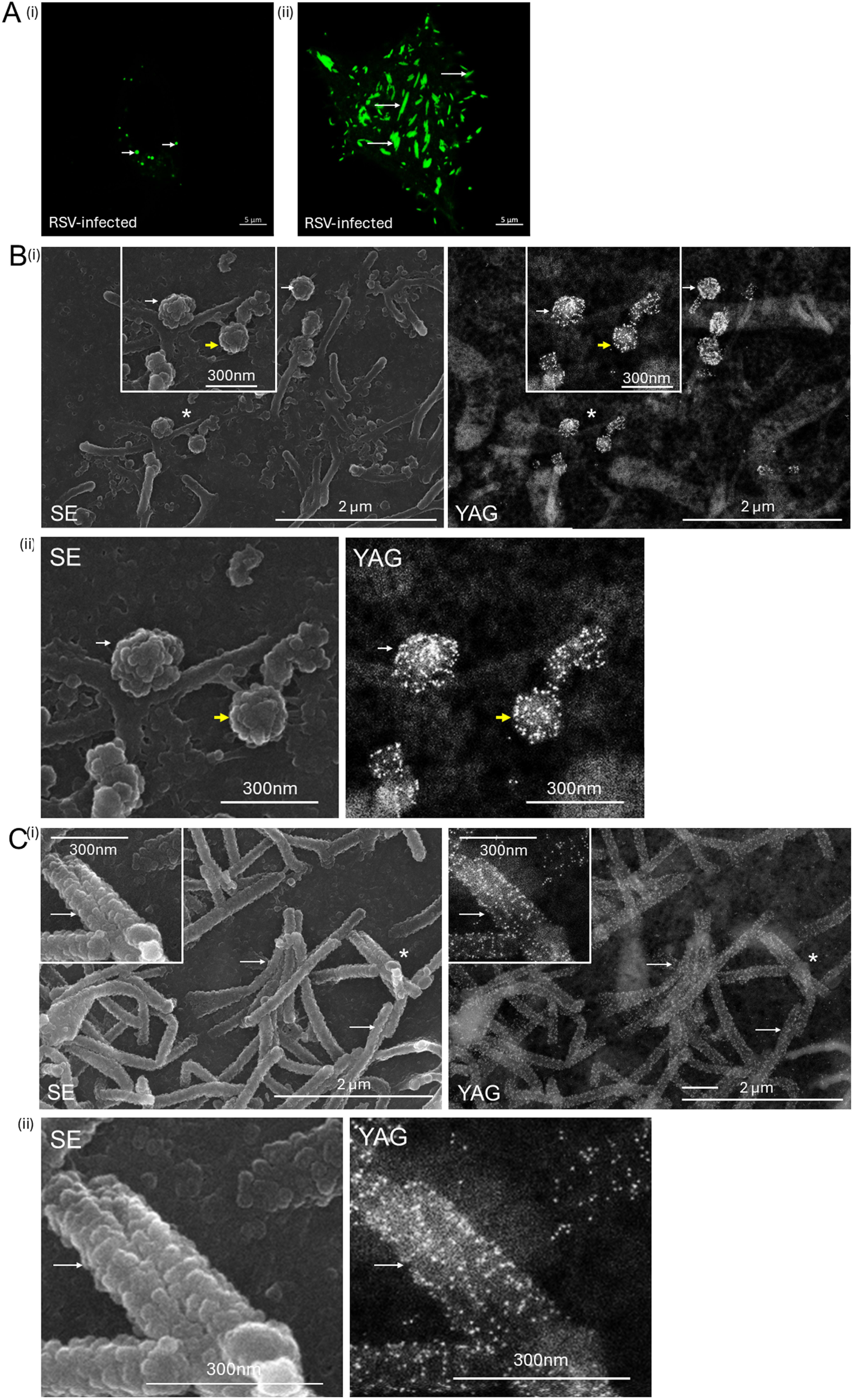
Imaging of the RSV particles on the surface of virus-infected cells at the early and late stage of infection. **(A)** A549 cells were infected with RSV and at (i) 2 hrs post-infection (hpi) and at (ii) 30 hpi the cells were incubated with anti-G and anti-mouse AL488 and imaged using confocal microscopy at a focal plane that allowed imaging of the cell surface (objective x100 magnification). The anti-G-stained virus particles at each time point are indicated (white arrows). **(B and C)** Cells were infected with RSV and at **(B)** 2 hpi and **(C)** 30 hpi the cells were incubated with anti-G and anti-mouse IgG conjugated to 10-nm colloidal gold and the surface of the monolayers was visualized in the FEG-SEM using the SE and YAG BSE (YAG) detector as indicated. The presence of gold particles in the YAG BSE image are presented as small bright points. In each plate (ii) is a further enlarged image taken from inset in (i) and the insets are obtained from the region demarcated (*) in (i). The spherical virus particles (short white arrows), the partially filamentous RSV particles (short yellow arrows) and the virus filaments (long white arrows) are highlighted.

The anti-G-labelled clusters were intermittent along the length of the virus filament, and although these clusters were not closely packed on the virus envelope, we estimated that any individual colloidal gold particle was within a radius of approximately 200-250nm to the next adjacent gold particle on the virus envelope (Fig. 2C(ii)). The limit of resolution of light microscopy is approximately 200nm, which together with the expected fluorescence dispersion, would account for the apparent continuous anti-G protein staining in the virus filaments that is detected using confocal microscopy as opposed to the intermittent anti-G staining revealed in the FEG-SEM analysis. Although the pleomorphic RSV particles detected on the cell surface at 2 hpi also exhibited a different particle morphology compared to the virus filaments, they exhibited a similar clustered anti-G staining pattern on their surface when compared with the virus filaments (Fig. 2B(ii)). This is consistent with the pleomorphic RSV particles originating from the virus filaments during the preparation of the virus inoculum. The analysis of the virus-infected cells using the SE detector at the later stages of infection (Fig 2C(i) and (ii)) again suggested a structured surface topology of the virus filaments and this was examined further using FEG-SEM and atomic force microscopy (AFM).

### Imaging the surface topology of virus filaments on the surface of virus infected cells using FEG-SEM and atomic force microscopy

#### 2.1 FEG-SEM

The antibody-labelled samples described to this point were coated with carbon, which is compatible with the use of the YAG BSE detector to localise colloidal gold conjugated antibodies (i.e. antigen localisation) on the surface of the virus filaments. However, a limitation is that a relatively thick carbon coating of 10-20 nm is required to achieve adequate conductivity, and that can mask fine surface detail on the surface of these structures. In a further analysis we therefore modified our existing protocols to improve the resolution of the surface topology of the virus filaments in the FEG-SEM analysis. The cells were coated with sputtered chromium (Cr) which has greater conductivity than carbon, allowing the application of thinner coatings over the samples (typically 2nm), and potentially higher resolution of the surface topology of the virus particles using the SE detector. In addition, chromium has spectral characteristics that are distinctly different from gold and the Cr coating can be used to visualise the location of the bound antibody-colloidal gold conjugates using the YAG BSE detector. To further improve electrical conductivity of the specimens, the cells were also cultured on silicon wafers rather than glass, since silicon is electrically conductive, thus enabling thinner coatings of Cr to be applied without electrostatically charging the biological specimens in the electron beam that would degrade imaging performance and resolution.

Initially RSV-infected cells were prepared for FEG-SEM and sputter coated with 2nm thickness of Cr and without antibody labelling. Examination of virus-infected cells at a lower magnification using the secondary electron (SE) detector revealed the presence of numerous virus filaments on the surface of the infected cells (Fig. 3A) that were similar in appearance to those described above in the carbon-coated specimens. The virus-infected cell surface was examined in more detail at higher magnification (Fig. 3B) to both examine the virus filaments and confirm the uniform 2nmCr coating over the specimen. An even and smooth Cr deposition over the entire specimen was noted, both on the bare silicon substrate where cells were absent and over which the virus filaments traversed (Fig. 3B (i) and (ii)), and across the surface of the infected cells where the virus filaments also formed (Fig. 3C)). This indicated the absence of any localised Cr deposition that would be expected to appear as visible clumps on the silicon substrate and cell surface. Examination of the cells at higher magnification showed that the surface of the virus filaments exhibited a structured appearance (Fig. 3B(ii)) and (Fig. 3C) that was distinct from the surface morphology exhibited on the cell surface, which suggested that the structured surface topology was a feature of the virus filaments and not of the Cr coating. Examination of these virus filaments at increasing magnification allowed their surface topology to be imaged in greater detail (Fig. 4) revealing a prominent cobblestone-like pattern, consisting of individual closely packed surface domains (Fig. 4A and B). These domains covered the surface of the virus envelope along the entire length of the virus filaments and were to be approximately 20-30 nm in diameter. This was distinct from the cell surface topology at locations where the virus filaments were absent, which appeared to have a relatively smoother texture when compared with the surface of the virus filaments (Fig. 4C). The glycocalyx coats the virus filaments and is expected to be visible on the surface of these structures, suggesting that the glycocalyx makes a major contribution to the distinct surface topology on the virus filaments that is revealed by the FEG-SEM. A similar structured cell surface topology has been reported on the cell surface of tissues when the glycocalyx was examined using scanning electron microscopy [36].

**Figure 3.**
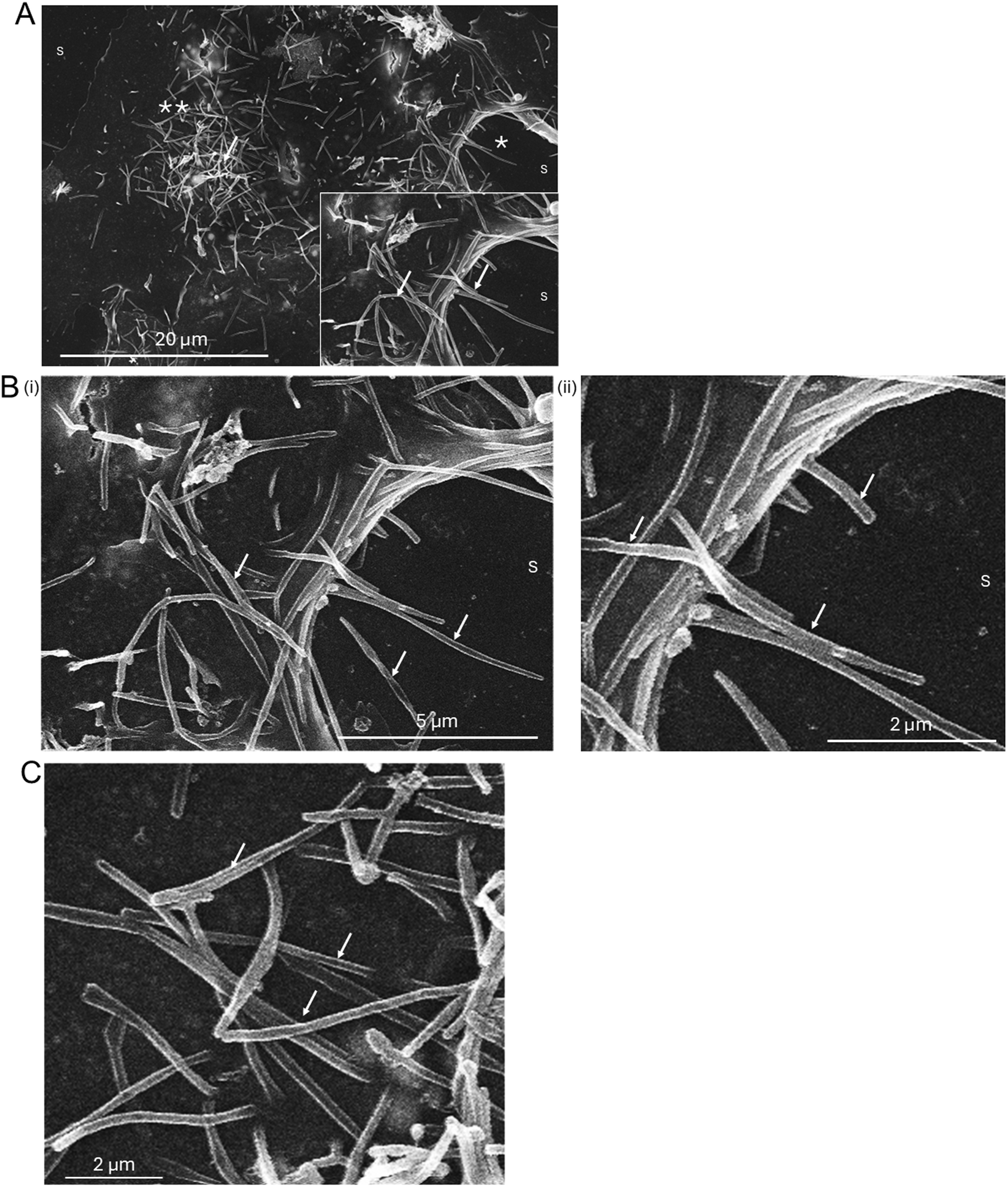
Imaging of the RSV particles on the surface of virus-infected cells after chromium coating. **(A)** At 20 hrs-post infection the RSV-infected A549 cells were coated with 2nm chromium and examined by FEG-SEM using the secondary electron detector (magnification x2,500). Inset is an enlarged image taken from the region (*) demarcated. The virus filaments (long white arrows) and the surface of the silica substrate (S) on which the cells are grown are highlighted. **(B)**(i) and (ii) and **(C)** are further enlargements of the areas in **(A)** that are demarcated by (*) and (**) respectively.

**Figure 4.**
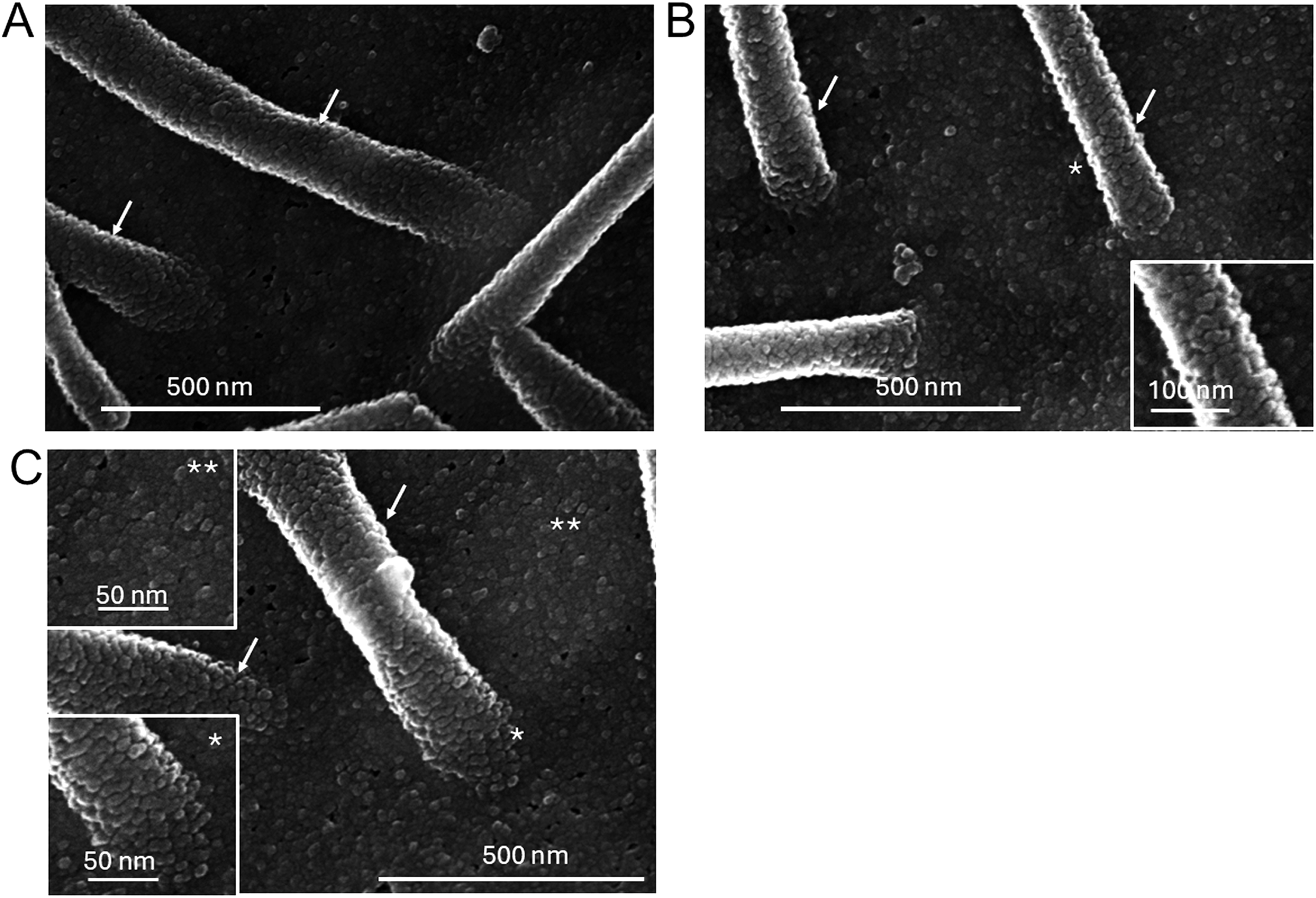
Imaging of the surface topology of RSV particles on the surface of virus-infected cells after chromium coating. At 20 hrs-post infection RSV-infected A549 cells were coated with 2nm chromium and examined by FEG-SEM using the secondary electron detector at **(A)** magnification x90,000, **(B)** magnification x100,000 and at **(C)** magnification x110,000. In each plate the virus filaments are highlighted (white arrows). In **(B)** the inset is an enlarged image taken from the region on the virus filament demarcated by (*) and in **(C)** the insets are enlarged images taken from the regions on the virus filament (*) and cell membrane (**) as indicated.

The identification of the virus filaments was confirmed by using ISEM in which virus-infected cells were labelled with anti-G and anti-mouse IgG-10nmAu prior to coating with the 2nmCr (Fig. 5). At lower magnification the presence of the virus filaments was defined using the SE detector (Fig. 5A(i) and B(i)), and the presence of anti-G staining along the lengths of the virus filaments was detected using the YAG BSE detector (Fig. 5A(ii) and B(ii))). A similar antibody staining pattern and staining distribution to that observed on the carbon-coated cells was noted, suggesting that the Cr coating did not affect the detection or the distribution of the anti-mouse IgG-10nmAu that was imaged by the YAG BSE detector. Although at many locations the anti-G staining again appeared to be clustered locally, the anti-G protein-stained clusters were intermittently distributed along the length of the virus filament. Examination of the virus filaments at even higher magnification (Fig. 5C(i)) and D(i)) again revealed the surface topology that appeared to be composed of individual interlocking surface domains that were similar to those observed on the unlabelled virus-infected cells as described above. In addition, the SE detector showed the presence of larger prominent globular structures (approximately 30-40nm in diameter) on the virus filaments whose appearance coincided with presence of the bound gold particles that were detected using the YAG backscatter detector (Fig. 5C(ii) and D(ii)). It was therefore concluded that these larger globular structures were due to the presence of the antibody conjugated to colloidal gold particles and identified the presence of the G protein in the virus envelope. This also suggested that a proportion of these individual interlocking surface domains that were observed on the surface of the virus filaments that form on the unlabelled virus-infected cells (i.e. in Fig. 4) contained localised concentrations of the G protein. However, anti-G staining was detected in a subset of these domains that were detected in the SE detector suggesting that these domains consist of concentrations of the G protein. In the ISEM analysis the antibody labelling is performed prior to sample preparation for electron microscopic imaging and location of the anti-G staining coincides with specific domains that are detected along the length of the virus filaments by the FEG-SEM. This suggests that the G protein distribution demarcated by the anti-G staining was not influenced by the subsequent sample preparation, further suggesting that the compartmentalisation of G protein on the virus envelope is a true reflection of the surface topology of the virus envelope. The other domains that remained unlabelled with anti-G and may consist of other factors that are displayed on the surface of the virus filaments, e.g. the host cell CD55 protein[12].

**Figure 5.**
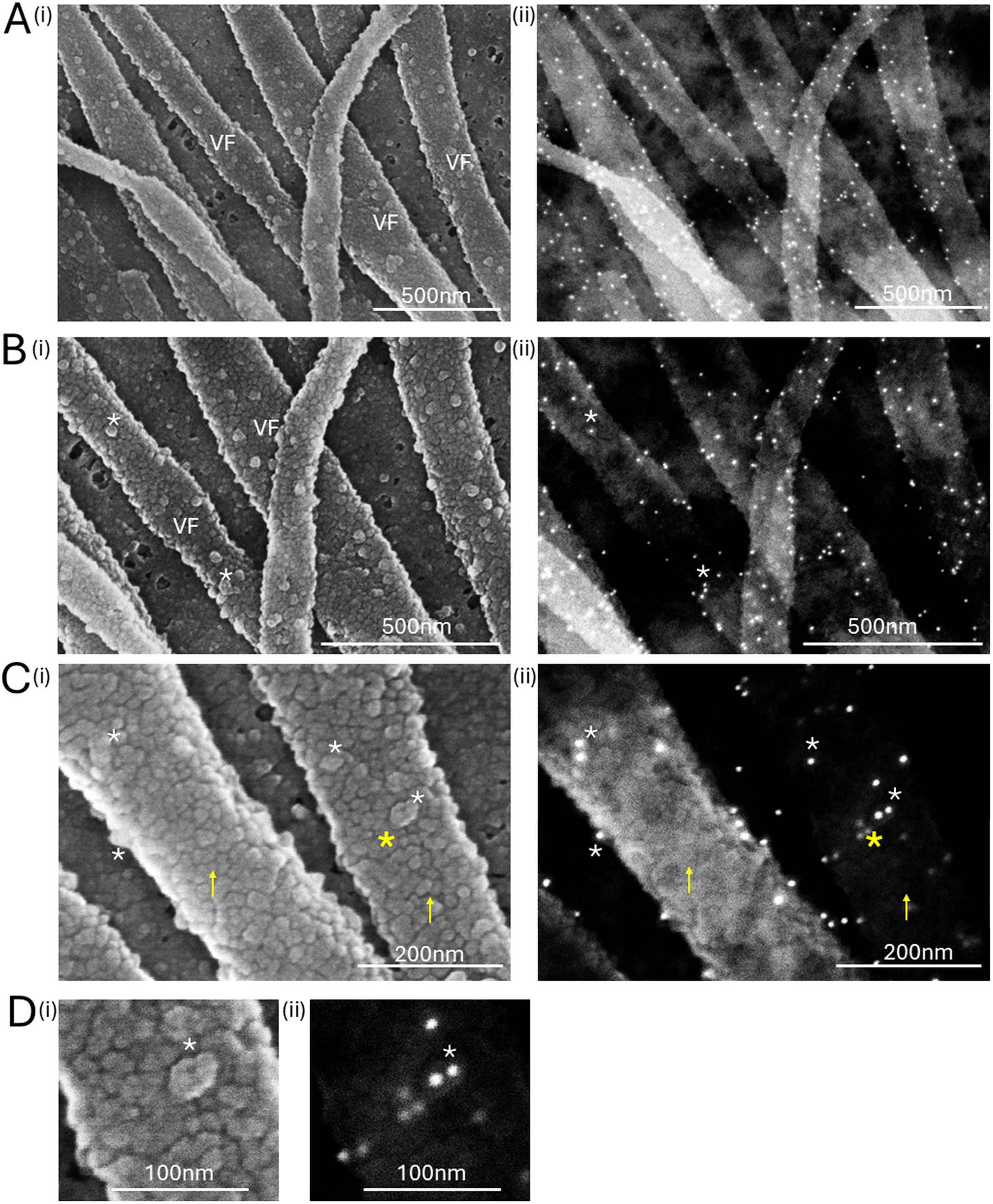
Imaging of the surface topology of RSV particles on the anti-G labelled virus-infected cells after chromium coating. At 20 hrs-post infection RSV-infected A549 cells were incubated with anti-G and anti-mouse IgG conjugated to 10-nm colloidal gold. The cells were coated with 2nm chromium and examined by FEG-SEM. The cells were examined at **(A)** magnification x70,000, **(B)** magnification x100,000 and **(C)** magnification x200,000 using the (i) using the secondary electron detector and (ii) YAG BSE detectors. The presence of the virus filaments (VF) is highlighted. The presence of the bound anti-G and 10-nm colloidal gold are highlighted (small white *) in SE and (ii) YAG BSE detector images, and the surface topology of the virus filament that is not bound with anti-G is also highlighted (yellow arrows) in the SE images. **(D)** is an enlarged image taken from the area indicated in **(C)** (highlighted by large yellow *) using the (i) SE and (ii) YAG BSE detectors. The presence of the bound anti-G and anti-mouse IgG conjugated to 10nm colloidal gold are highlighted (small white *) in (i) SE and (ii) YAG BSE images,

The analyses described above using the FEG-SEM on the Cr-coated cells were performed using a standard accelerating voltage of 10kv, which gives good resolution in the SE mode in the FEG-SEM and good signal in the BSE mode. The FEG-SEM analysis on the Cr-coated anti-G labelled virus-infected cells was also performed using a lower accelerating voltage of 5kv. Under these imaging conditions the electron beam is less penetrating while high resolution is still maintained enabling the surface topology of the virus filaments to be imaged, and imaging of the virus filaments with the SE and YAG BSE detectors was performed at this reduced accelerating voltage (Fig. 6). Although there was a reduced signal emission from the sample in both the SE and YAG BSE detectors when compared with that image using 10 kV, the recorded images showed that the surface of the virus filaments exhibited a similar overall surface topology and anti-G staining distribution on to that imaged using the higher accelerating voltage.

**Figure 6.**
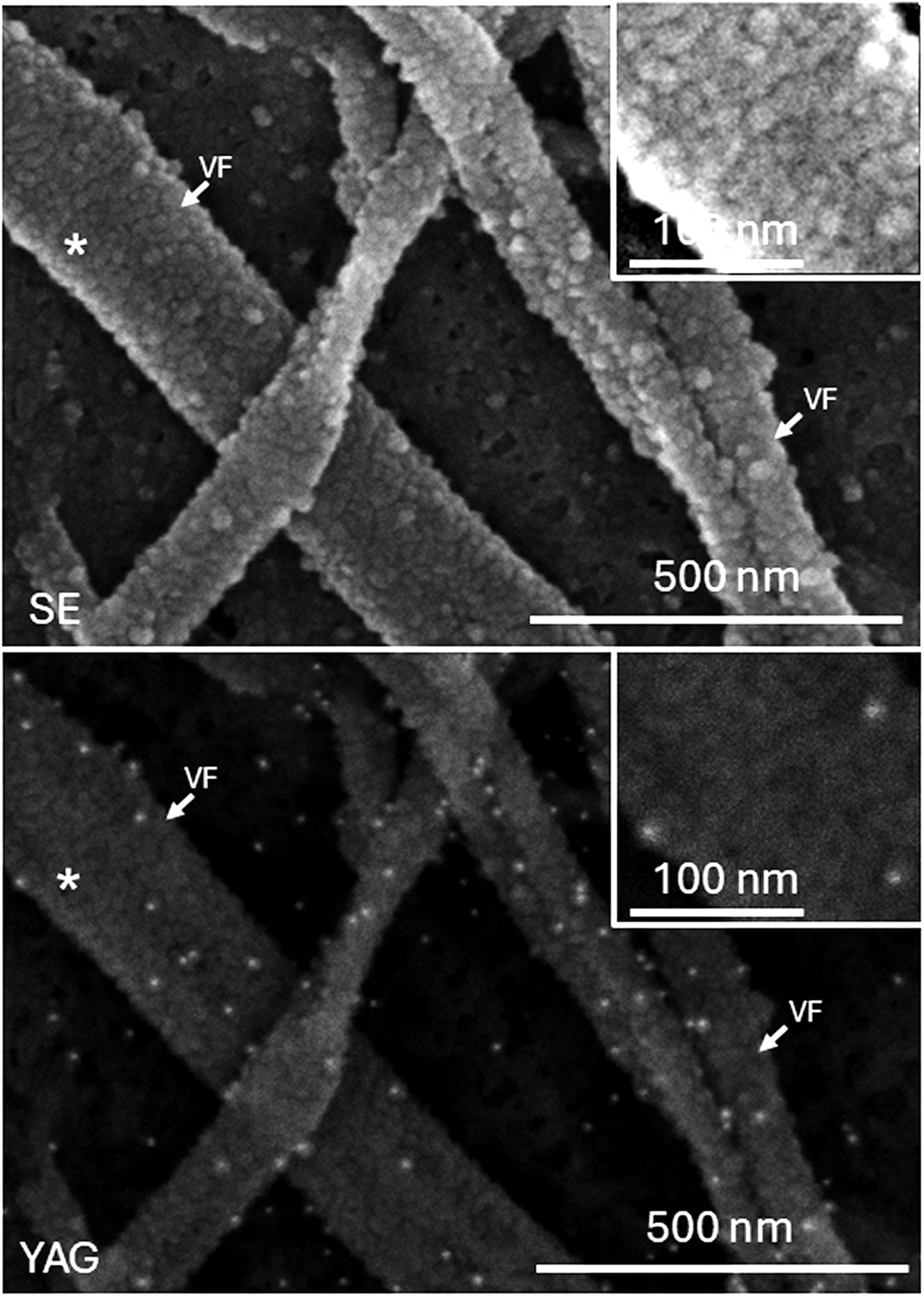
Imaging of the surface topology of chromium coated-RSV particles on the anti-G labelled virus-infected cells using reduced accelerating voltage. At 20 hrs-post infection RSV-infected A549 cells were incubated with anti-G and anti-mouse IgG conjugated to 10-nm colloidal gold. The cells were coated with 2nm chromium and examined by FEG-SEM. The cells were examined using the FEG-SEM at magnification x100,000 at an accelerating voltage of 5 kV using the secondary electron (SE) and YAG BSE (YAG) detectors as indicated. The virus filaments (VF) are highlighted and the presence of gold particles in the YAG BSE image are presented as small bright points. The insets are enlarged images of the virus filaments taken from the area that is highlighted (*) in the main plate.

The immunolabelling in the ISEM analysis that was described above was performed using anti-mouse IgG (whole molecule) conjugated to 10nm colloidal gold to detect the presence of the bound anti-G antibody. Although in the YAG BSE detector it is the 10nm gold particle that is detected, a significant contribution to the particle size (i.e. the molecular volume) of the antibody conjugated gold particle comes from the antibody that is used to coat the colloidal gold particle [37]. This can be seen at the locations of bound anti-mouse IgG-10nmAu in the images obtained in the SE detector (Fig. 5E(i)) and the YAG BSE detector (Fig. 5E(ii)), where the difference in size of the signal due to the 10nm gold particle (visualised in the YAG-image) and the anti-mouse IgG-10nmAu (visualised in the SE-image) can be compared. We therefore also performed this analysis using antibodies conjugated to a smaller colloidal gold particle of 5 nm (5nmAu), to ensure that the antibody distribution that we observe was not due to the use anti-mouse IgG-10nmAu. In this analysis the cells were labelled with anti-G and then either anti-mouse IgG-5nmAu (Fig. 7A) or anti-mouse Fab-5nmAu (Fig. 7B), the latter being an even smaller antibody probe consisting of only the antibody Fab fragment conjugated to 5nm colloidal gold. The anti-mouse Fab-5nmAu has a smaller molecular volume when compared with the colloidal gold particle conjugated to the whole IgG. The labelled cells were coated with the 2nm Cr and examined by FEG-SEM using both the SE detector and YAG BSE detector. The SE detector showed the presence of the virus filaments which coincided with the bound gold particles that were visible using the YAG BSE detector (Fig. 7A(i) and 7B(i)). Examination of the immunolabelled virus filaments at higher magnification using the SE detector showed the individual interlocking surface domains structures on the surface of the virus filaments and the YAG BSE detector revealed the presence of the anti-G staining on a proportion of these structures (Fig. 7A(ii) and 7B(ii)). A slightly higher number of gold particles per G protein cluster was noted when labelling was performed with the anti-mouse Fab-5nmAu probe when compared with the larger anti-mouse IgG-5nmAu. This may indicate that the smaller gold probe was able to recognise more of the anti-G bound to the G protein clusters and suggests that there may be a degree of steric hinderance when the whole IgG molecule is used that effects the stoichiometry of the secondary antibody binding to the G protein clusters. However, a similar surface topology and a similar intermittent clustered anti-G staining pattern was observed for the secondary antibodies conjugated to the 5nm colloidal gold (IgG or Fab) and 10nm colloidal gold. The intermittent anti-G staining pattern was therefore not related to the size of the colloidal gold conjugate used in the immunolabelling and reflects the distribution of the G protein clusters on the surface of the virus filaments.

**Figure 7.**
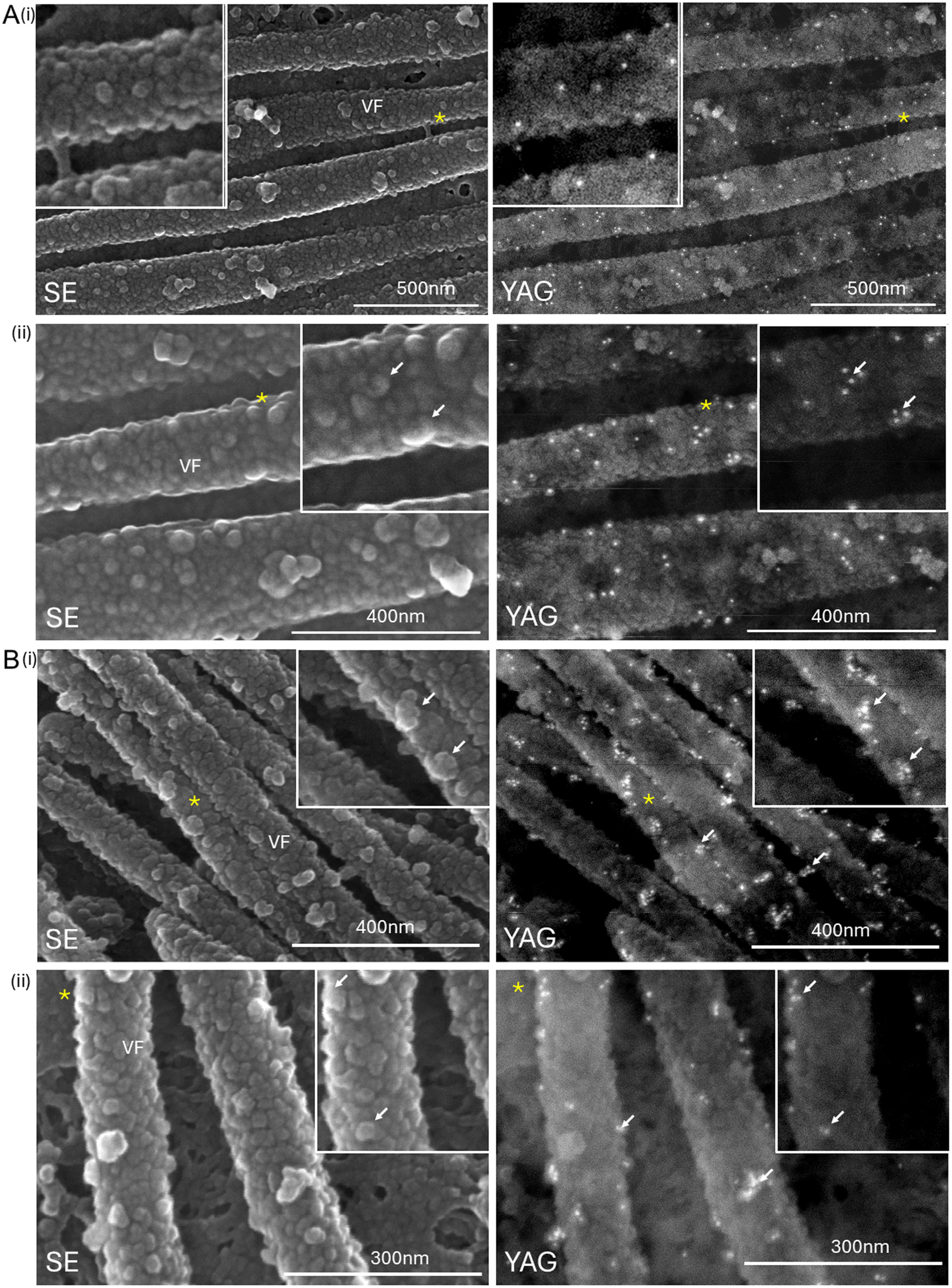
Distribution of the G protein on the surface of RSV particles using smaller gold probes. At 20 hrs-post infection RSV-infected A549 cells were incubated with anti-G and **(A)** anti-mouse IgG conjugated to 5 nm colloidal gold or **(B)** anti-mouse Fab conjugated to 5nm colloidal gold. The cells coated with 2nm chromium and the surface of the monolayers was visualized in the FEG-SEM using the secondary electron (SE) and YAG BSE (YAG) detector as indicated. Imaging was performed at **(A)** (i) magnification x70,000, and (ii) magnification x130,000 and (**B)** (i) magnification x80,000 and (ii) magnification x130,000. The virus filaments (VF) and the presence of gold particles in all YAG images (white arrows) are highlighted. In all plates the insets are enlarged images taken from the regions highlighted (yellow *) in the main plate.

#### 2.2 Atomic force microscopy

Atomic force microscopy (AFM) was also used to image the surface topology of the virus filaments that form on RSV-infected cells. Although the resolution of AFM is expected to be lower than FEG-SEM analysis that was described above, it allowed the surface topology of the RSV filaments to be examined without the requirement of a conductive coating and the cells remain in a hydrated state. It therefore represents a useful complementary approach with which to compare the surface topology of the virus filaments that were imaged using FEG-SEM procedure. The AFM analysis uses three different machine configurations to image the surface topology of the virus filaments. The hight-mode (H-mode) and amplitude-mode (A-mode) refer to the height of the virus filaments relative to the substrate on which the cells are attached. The phase-mode (P-mode) is an additional configuration that captures information based on the material properties of the specimen under examination (e.g. the virus filaments) and this provides additional topological information related to the virus filaments.

Cells were infected with RSV and at 20 hpi the cells were examined by AFM to determine the surface topology of the virus filaments. Examination of the RSV-infected cells by AFM using a wide scan size of 10 μm in A-mode and H-mode allowed single cells to be examined, and this showed the presence of numerous long filamentous structures that were greater than 1 μm in length on the cell surface (Fig. 8A). These filamentous structures exhibited physical dimensions that are similar to the virus filaments that were observed on virus-infected cells using the FEG-SEM at low magnification. These virus filaments were examined in greater detail in A-mode and H-mode using a narrow scan size of 1.5 μm (Fig. 8B). The surface topology of the virus filaments exhibited a similar structured appearance and exhibited a similar overall appearance to that observed by the FEG-SEM analysis. This was more clearly demonstrated when the data obtained in the A-mode were rendered into a 3-dimensional image where the virus filaments that were imaged extended across the cell surface (Fig. 8C and D). The surface topology of the virus filaments that form on the surface of the virus-infected cells were examined in even greater detail using a lower scan size (Fig. 9) which was equivalent to the higher magnifications used in the FEG-SEM analysis. This was performed in A-mode, H-mode and P-mode on two representative clusters of virus filaments that formed on the surface of infected cells, using a scan size of 666.5 nm (Fig. 9A), and 603.9 nm and 302nm (Fig. 9B). In each virus filament cluster, a similar representation of the surface topology on the virus filaments was noted for each of the three modes used. This revealed the presence of a structured surface topology on the surface of the virus filaments that consisted of individual closely packed surface domains that was similar in appearance to the surface topology imaged using FEG-SEM. The domains detected in the AFM analysis had an average diameter of approximately 30-40nm and were slightly larger than those detected in the FEG-SEM analysis. This may be partly due to the differences in the methodologies associated with the sample preparations and the ways in which the different methods acquired the images of the virus filaments. For example, in AFM the cells are not coated with a conductive material, and they are maintained in a hydrated state, and while the SEM detects the presence of scattered electrons emitted from the specimen in the electron beam, the AFM measures amplitude changes in the sample by micro cantilever probe displacement. However, irrespective of these small size differences a similar structured surface topology was observed on the virus filaments using the AFM and FEG SEM procedures, and this consistency suggested that the surface topology of the virus filaments consisted of ordered interlocking domains.

**Figure 8.**
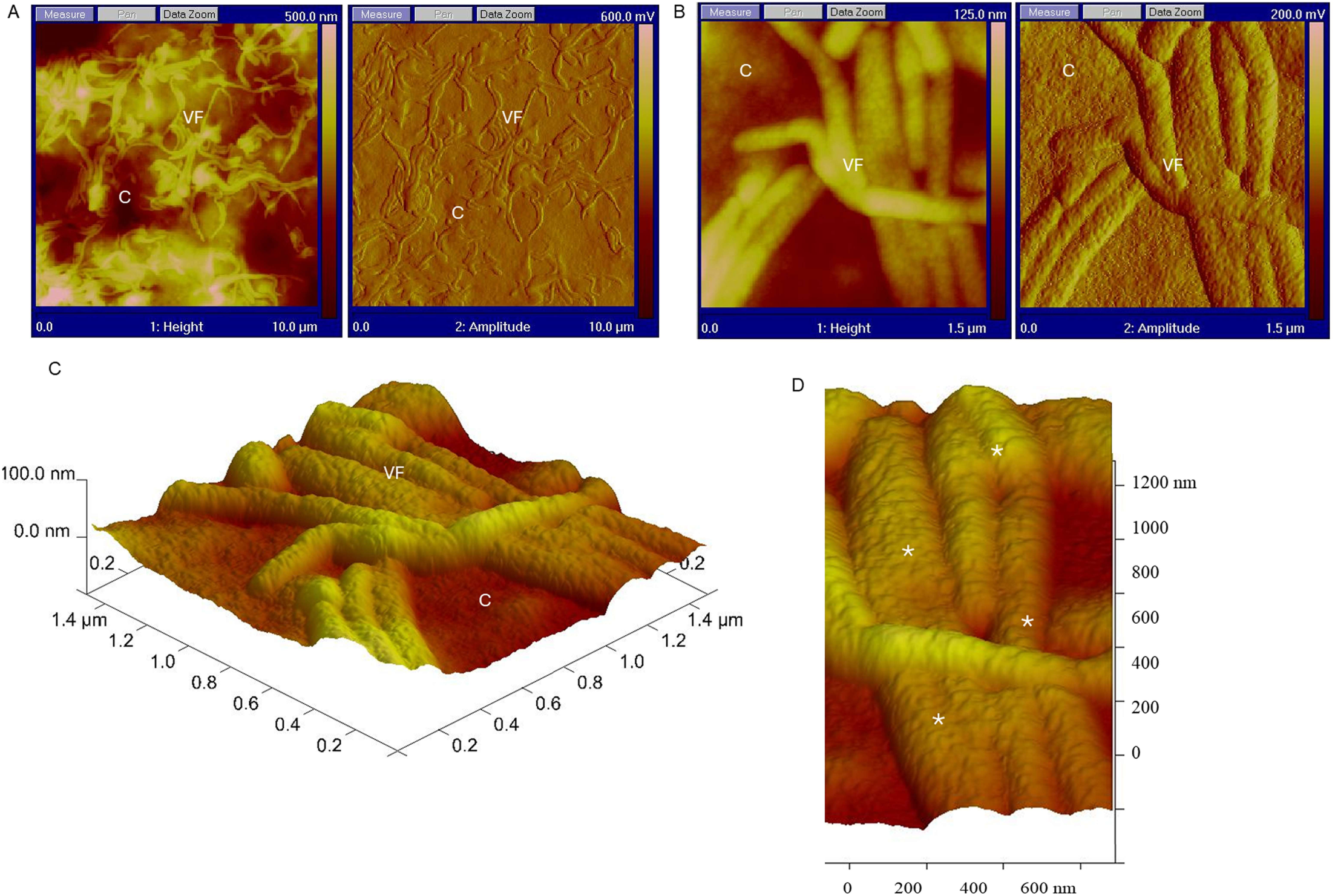
Imaging of the RSV particles on the surface of virus-infected using atomic force microscopy (AFM). At 20 hrs-post infection RSV-infected A549 cells were examined using AFM in height and amplitude mode as indicated using a scan size of **(A)** 10μm and **(B)** 1.5μm. (**C**) The data obtained using the amplitude mode was rendered into a 3-dimensional image and (**D**) is an enlarged image from the 3-dimensional image in **(C)**. The virus filaments (VF), cell surface (c) and the topology on the virus filaments (*) are highlighted.

**Figure 9.**
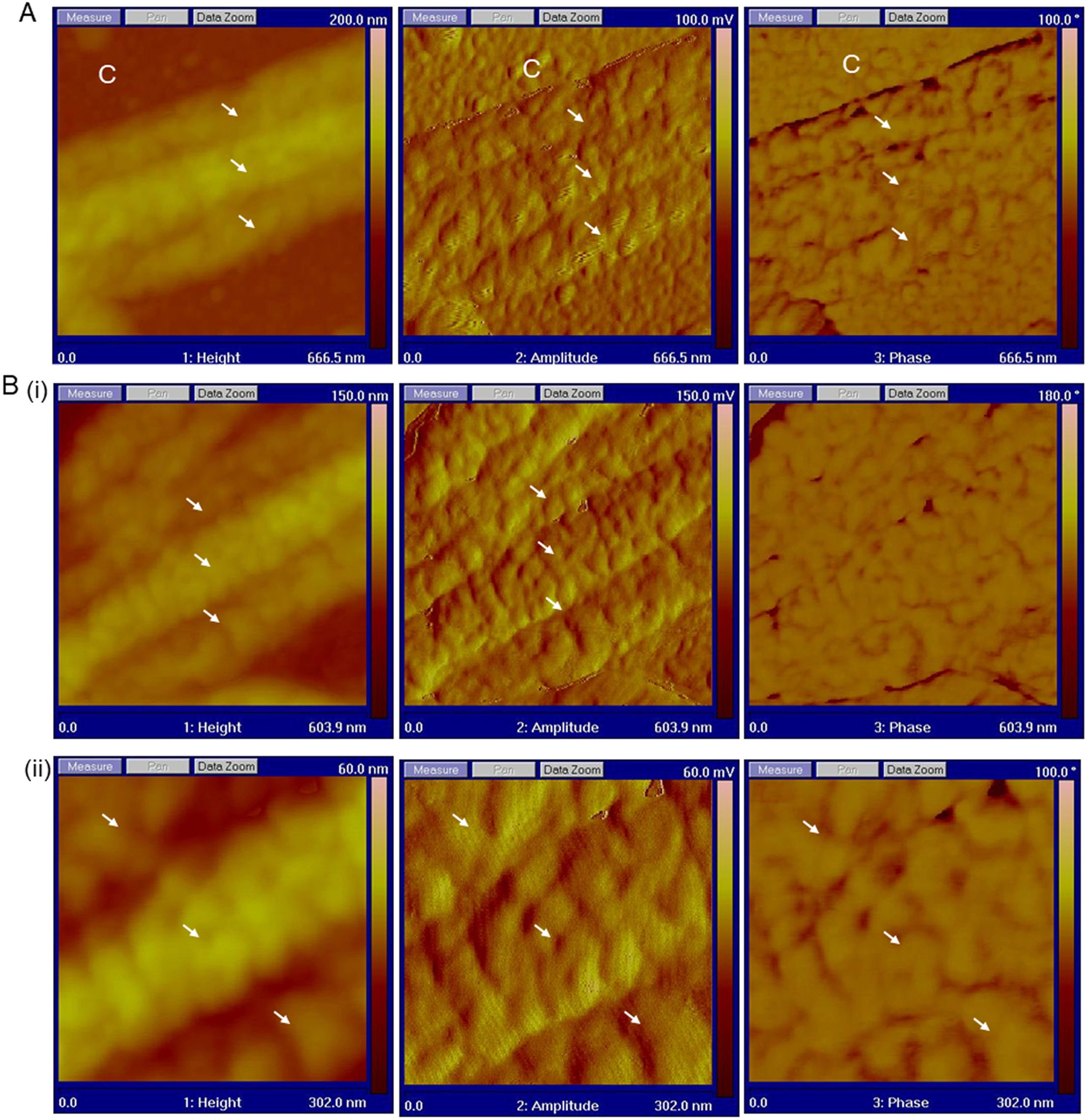
Imaging of the surface topology of RSV particles on the virus-infected cells using atomic force microscopy (AFM). Two cluster of virus filaments **(A)** and **(B)** on the surface of RSV-infected A549 cells were examined at 20 hrs-post infection using AFM in height, amplitude and phase mode as indicated, using a scan size of **(A)** 666.5 nm and **(B)** (i) 603.9nm and (ii) 302nm. The surface topology on the virus filaments is highlighted (white arrows).

### 3. Conclusions

In this study we have performed a high resolution *in situ* image analysis of the surface topology of virus filaments as they formed on the surface of RSV-infected cells. We provide evidence for a compartmentalised and ordered surface topology consisting of individual domains that are closely packed along the length of the virus filaments. This is perhaps not surprising since a variety of different host cell factors have been detected on the surface of these structures. This includes cellular protein networks that regulate the complement cascade [12, 38] and proteins that form the cell glycocalyx that envelope the virus filaments [31]. Our current study using FEM-SEM is also consistent with previous analysis of virus filaments using confocal microscopy imaging. These previous analyses demonstrated that the virus-associated cellular factors that were identified exhibited a staining pattern within the virus filaments that was consistent with their localised concentration along these virus filaments. The compartmentalisation of these cell factors on the virus filaments may be an important factor for the functionality of these cellular factors within the virus envelope. The G protein is expressed in high abundance on the surface of virus filaments with respect to other virus proteins that are displayed on the surface of these structures. Our current analysis suggests that the G protein was also clustered within specific domains on the virus envelope that appear to be intermittently arranged along the length of the virus filaments. The localised clustering of the G protein in the virus envelope is expected to increase the concentration of the G protein at these sites which may in turn have important implications for the way that the G protein mediates the interaction of the virus with the surface of the host cell during the early stages of virus entry. Although the glycocalyx coats the entire cell surface, the glycocalyx appears to be more highly concentrated in cell surface projections such as the microvilli, suggesting that these structures may have increased levels of the glycocalyx [31]. The recombinant expressed G protein is transported into these structures, and furthermore the G protein is also located within the glycocalyx that surrounds the virus filaments on virus-infected cells [31]. These observations suggest that RSV uses preexisting cellular structures on the cell surface as the site of virus particle assembly, and that the glycocalyx that is present on these structures may also contribute to the structured surface topology of the virus filaments that we observe in the current imaging analysis. Finally, this study emphasises the utility of using ISEM as experimental approach to further characterise the organisation of the surface topology of the virus particles on the surface of virus-infected cells, and this will be a focus of future work. Using ISEM to examine RSV assembly *in situ* on cells will serve as complementary experimental approach to the high-resolution imaging techniques that are used to examine RSV ultrastructure.

## Materials and methods

### Cells and viruses

The RSV A2 strain was used throughout this study and prepared as described previously [32]. A459 cells were maintained in growth medium; Dulbecco’s Modified Eagle’s Medium (DMEM) supplemented with 10% foetal calf serum (FCS) and antibiotics.

### Virus infection

The A549 cells were infected with RSV at 33°C and 5%CO_2_ in a humidified chamber using a multiplicity of infection of 5. At 2 hrs post-absorption the inoculum was removed and the cells washed and incubated using maintenance media (DMEM+2%FCS and antibiotics). Mock-infected cells were prepared in a similar manner using maintenance media instead of virus inoculum. The mock-infected and virus-infected cells were continued in maintenance media until the time of harvesting for down-stream processing.

### Specific reagents

The RSV antibody that recognises the G protein (MAb30) protein were provided by Geraldine Taylor (IAH,UK), and the anti-SH antibody has been described previously[35]. The anti-mouse AL488 (invitrogen) and the anti-mouse IgG-10nmAu, anti-mouse IgG-5nmAu and anti-mouse Fab-5nmAu (British BioCell International) were purchased. The silicon wafers were purchased (TAAB) and washed using absolute ethanol, sterile distilled water, followed by growth medium before use.

### Immunofluorescence microscopy

Cells were fixed with 3% (w/v) paraformaldehyde in PBS for 20 minutes at 25°C and washed extensively with PBS. The cells were incubated with the MAb30, and the cells were washed and incubated for a further 1 hour with the secondary antibody (goat anti-mouse IgG (whole molecule) conjugated to Alexa 488 (Molecular probes) and the stained cells were mounted on glass slides using Citifluor. The stained cells were visualised either by immunofluorescence microscopy using a Nikon Eclipse 80i Microscope (Nikon Corporation, Tokyo, Japan) and images processed using Q Capture Pro ver. 5.0.1.26 (Q Imaging, Teledyne Photometrics) or by using a Zeiss 710 confocal microscope with Airyscan using appropriate machine settings and the images processed using Zen ver 2.3 software.

### Immuno-scanning electron microscopy

Cells seeded on glass coverslips or silica wafers were fixed with 0.1% glutaraldehyde in PBS at 4 °C for 30 minutes. The fixed monolayers were incubated in PBS supplemented with 2 mM lysine for 20 minutes after which they were washed extensively with PBS. The cells were incubated at 25°C for 4 hour with MAb 30 or anti-SH and then incubated for a further 4 hours in goat anti-mouse IgG (whole molecule) conjugated to either 5 or 10 nm colloidal gold or IgG (Fab) conjugated to 5 nm colloidal gold. The monolayers were extensively washed with PBS and fixed with 2.5% glutaraldehyde in PBS and processed for SEM as described previously [2]. The samples were sputtered with carbon or chromium and visualised in a Hitachi 4700F field emission scanning electron microscope using appropriate settings. Digital images were recorded using Hitachi FE PCSEM (ver 3.2) software.

### Atomic Force Microscopy (AFM)

Atomic Force Microscopy was performed on infected cells on silica wafers using a Nanoscope IV MultiMode AFM (Veeco Instruments). Imaging was performed in dry Tapping Mode using DNP-probes using a scan rate of 1Hz and 256 samples per line and 256 lines using line direction in retrace mode. The data was processed using software Nanoscope version 6.11r1 (Veeco).

## ACKNOWLEDGEMENTS

We thank Jim Aitken and Gaie Brown for technical support. This work was supported by the MRC (UK) and the MOE Singapore.

